# Inhibition of Ras1-MAPK pathways for hypha formation by novel drug candidates in *Candida albicans*

**DOI:** 10.1101/2021.07.06.451239

**Authors:** Young Kwang Park, Jisoo Shin, Hee-Yoon Lee, Hag Dong Kim, Joon Kim

## Abstract

The opportunistic human fungal pathogen *Candida albicans* has morphogenesis as a virulence factor. The morphogenesis of C. albicans is closely related to pathogenicity (1). Ras1 in *C. albicans* is an important switch in the MAPK pathway for morphogenesis (2, 3). The MAPK pathway is important for the virulence, such as cell growth, morphogenesis, and biofilm formation (4, 5). Ume6 is a well-known transcriptional factor for hyphal-specific genes (6). Despite numerous studies, as a recent issue, it is necessary to develop a new drug that uses a different pathway mechanism to inhibit resistant *C. albicans* strains caused by chronic prescription of azole or echinocandin drugs, which are mainly used. Here, we show that the small carbazole derivatives attenuated the pathogenicity of *C. albicans* through inhibition of the Ras1/MAPK pathway. We found that the small molecules inhibit morphogenesis through repressing protein and RNA levels in Ras/MAPK related genes including *UME6* and *NRG1*. Furthermore, we found the antifungal effect of the small molecules *in vivo* using a candidiasis murine model. We anticipate our findings are that the small molecules are the promising compounds for the development of new antifungal agents for the treatment of systemic candidiasis and possibly for other fungal diseases.

**Author summary:** The infection by the opportunistic human fungal pathogen *Candida albicans* occurs mainly in immunocompetent and immunocompromised humans, such as AIDS patients, immunosuppressant-treated organ transplant patients, and recent COVID-19 patients. Morphogenesis which the ability to switch between yeast and hyphal growth forms is one of the representative virulence factors of *C. albicans.* Here, we describe novel small molecules that show antifungal effects such as the inhibition of the morphogenesis and the biofilm formation, and maintenance of biofilm. Moreover, we found that these small molecules had antifungal activity in mouse experiments, and confirmed that they were also effective in drug-resistant *C. albicans* strains. Studies of some small molecules with structures similar to ours have already been reported to exhibit growth inhibitory activity against bacteria and *Candida* species. However, the mechanism of action of these molecules has not been elucidated. In this study, we demonstrated, for the first time, the mechanism by which these two small molecules inhibit *C. albicans* pathogenicity through inhibition of specific pathways. Our study, through the research of the mechanism of action of novel small molecules, provides new insights into the development of drug candidates not only for wild-type *C. albicans*, but also for strains resistant to existing drugs.

## Introduction

*Candida albicans* causes opportunistic infections in humans as a commensal fungal pathogen. Life-threatening systemic fungal infections occur mainly in immunocompetent and immunocompromised humans, such as AIDS patients, immunosuppressant-treated organ transplant patients, and recent COVID-19 patients (7). Candida infection has shown an increasing prevalence and severity with an increase in AIDS, chemotherapy, and organ transplantation. Candidemia accounts for about 9% of nosocomial bloodstream infections, and when *C. albicans* invades the internal organs, it can be severe candidiasis with a mortality rate of approximately 40%.

Morphogenesis is one of the representative virulence factors of *C. albicans* (8). It has two main forms: yeast and hyphal. The dimorphism in which the ability to switch between yeast and hyphal growth forms is essential for pathogenicity. The yeast form is required for adhesion and colonization of the epithelial cells of the host, while the hyphal form is required for the penetration and invasion of endothelial cells of the host(9). Morphogenesis has various environmental cues, such as nutrient limitation, alkaline pH, quorum sensing molecules, serum, elevated temperature, elevated CO_2_, and embedded conditions (3, 10). For *in vitro* experiments, elevated temperature to 37°C with serum was used for critical hyphal formation stimulation (11). Importantly, morphogenesis is linked to the virulence of *C. albicans* (12). Therefore, the general screening for novel drugs for *C. albicans* commonly uses the inhibition of morphogenesis as a benchmark.

In *C. albicans*, MAPK and Ras/PKA-dependent pathways govern morphogenesis as well as nutrient sensing and acquisition, stress response, and pathogenesis. These pathways play a potent role in hyphal formation in *C. albicans*. Various transcriptional factors have been identified as signal transduction regulators in these pathways. The small GTPase Ras1 in this fungus affects both pathways as a switch for hyphal formation signals, such as serum, elevated temperature, and nutrient limitation. As one of the MAPKs in *C. albicans*, Cek1 has a role in biofilm formation and filamentous growth in *C. albicans*, while another MAPK, Mkc1, has a function in the cell wall structure and maintenance. The representative transcription factor that mainly influences hyphal formation is CaUme6, which upregulates hyphal-specific genes (HSGs) encoding hyphal cell-related components such as Hwp1, Als3, and Ece1 proteins. Transcription factors such as Ume6, Efg1, Cph1, and Cph2 are also involved in hyphal elongation; notably, Ume6 plays a role in filamentous growth maintenance (6). Overexpression of CaUme6 induced distinct hyphal formation, whereas downregulated CaUme6 could not induce hyphal formation even at elevated temperatures (37°C) with serum incubation conditions. The final recipient of many hyphal growth signaling pathways in *C. albicans* is Ume6 and as a key controller in hyphal growth and the transcriptional repressor, Nrg1 works with Ume6 as a negative feedback loop to control the expression of HSGs in hyphal growth environmental signals.

Fluconazole, an azole drug typically used as an antifungal agent, targets the enzyme essential for cell wall ergosterol biosynthesis. As another generally used antifungal drug, amphotericin B (polyene) and echinocandin target fungal membrane and cell wall synthesis respectively. In the ergosterol biosynthesis pathway, azoles inhibit ERG11, which encodes 14α-demethylase (cytochrome P450 enzyme lanosterol demethylase) (13). The azole-resistant *C. albicans* strains have been isolated from chronically prescribed chemotherapy patients. The resistant *C. albicans* strains have some features such as point mutations and the overexpression of ERG11, the azole target, and the overexpression of drug efflux pumps Cdr1 and Mdr1. To overcome these azole-resistant *C. albicans* strains, the necessity of new antifungal drugs that have different targets from those commonly used is emphasized.

Here, we describe novel small molecules that inhibit morphogenesis at elevated temperatures with serum incubation under hyphal induction conditions. These small molecules show antifungal effects at MIC and appear to inhibit the formation and maintenance of biofilm, the ultimate form of hyphal morphogenesis. Through a murine candidiasis model, these small molecules have been shown to have potential as new antifungal drugs *in vivo*. We identified the inhibition of Ras1-Cek1 signal transduction by these small molecules via the transcriptional and translational levels of each pathway regulator.

## Results

### Identification of small molecules that inhibit the growth of *Candida albicans*

In our previous study (14), it was confirmed that the vacuole inhibitors have antifungal activity, and 32 relatively effective analogs of piperazinyl carbazole structure which was anticipated to possess vacuole inhibitory activity of bafilomycin A1 and compounds containing the carbazole structure(15–17) were also considered and screened. The active small molecules consist of carbazole, piperazine, and phenyl amine groups (Fig. 1A). To screen for small molecules that have growth inhibitory effects on *C. albicans,* MICs of small molecules were investigated according to the Clinical and Laboratory Standards Institute (CLSI) document M27-A3 (18), resulting in identification of two compounds showing inhibitory effects on the growth of *C. albicans* (Fig. 1B and S1A). As shown in Table 3, the MICs of these small molecules were 16 μg/mL and 8 μg/mL, respectively, in SC5314, the wild type strain of *C. albicans.* Moreover, these two small molecules both showed MICs in BY4741, the wild type strain of *S. cerevisiae* as 4 μg/mL (Fig. S1B). Furthermore, these molecules showed noncytotoxicity in mammalian cells at MIC (Fig. 1C and S1C).

**Fig. 1.**
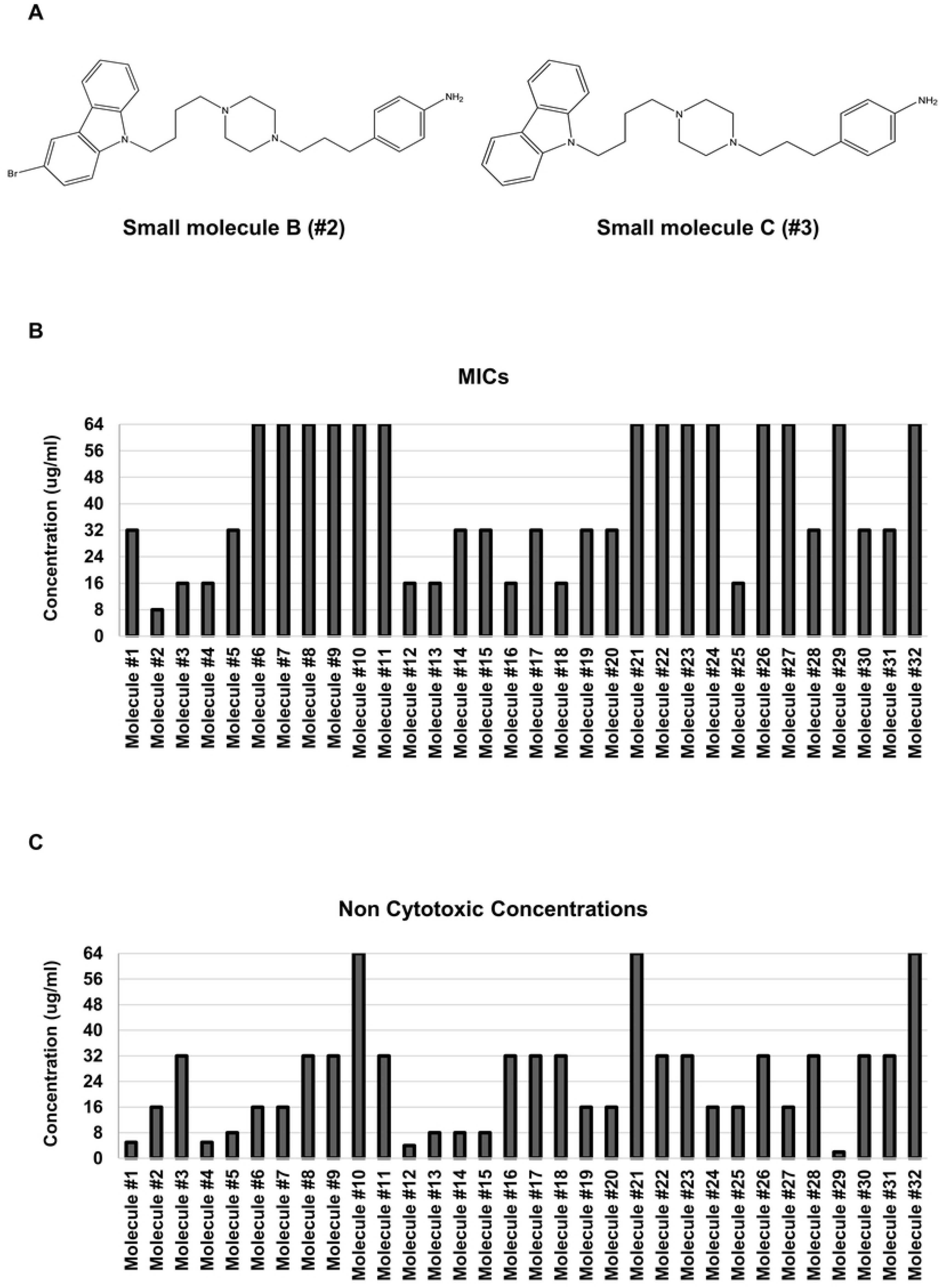
The cytotoxicity of small molecules in *C. albicans* and mammalian cell, and the chemical structures of the small molecules. (A) Structures of the small molecules B(#2) and C(#3). (B) Minimum inhibitory concentrations (MICs) of small molecules. *C. albicans* cells (2 × 10^4^ cells/mL) were grown with the small molecules in RPMI1640 medium with 0.165 M MOPS for 24 h at 37°C according to CLSI guidelines (18). The growth inhibition was determined by measuring the optical density at 595 nm using a microplate reader. MICs were defined as the lowest concentration of the small molecules that inhibited the growth of the cells by 95% compared with the control. (C) Analysis of the cytotoxicity of the small molecules on HeLa cells. The cytotoxicity was determined using the MTS assay. The noncytotoxic concentrations were defined as the highest concentration of the small molecules that inhibited the growth of the cells by 5% compared with the control. The absorbance at 490 nm was measured with a microtiter plate reader. Each experiment was conducted in triplicate.

Surprisingly, the MICs for fluconazole-resistant and echinocandin-resistant *C. albicans* strains of these small molecules showed the same MIC as the wild type strain (Table 3). These results suggest that the small molecules have growth inhibitory activity against *C. albicans,* including strains that are resistant to existing drugs.

### Inhibitory effects of the small molecules on morphogenesis and biofilm formation in *Candida albicans*

Growth inhibition is insufficient to prevent the virulence of *C. albicans.* Morphogenesis, a notorious virulence factor in *C. albicans*, is used as the standard for general testing of new drugs against *C. albicans*. To investigate the effect of inhibiting the morphogenesis of small molecules, hyphal formation was induced according to the presence or absence of small molecules. To induce hyphal morphogenesis, a YPD medium containing 10% FBS at 37°C was used and a YPD medium at 30°C incubation was used for the yeast form control. Small molecules B and C were used to treat hyphae formation at the concentration of each MIC. As a control group, the yeast form was shown in YPD medium at 30°C incubation, and the hyphal form was shown in YPD medium containing 10% FBS at 37 °C. Small molecules B and C completely inhibited hyphal formation under the conditions that stimulated hyphal morphogenesis (Fig. 2A and 2B). Biofilm, a structured microbial community, is the final form of hyphal development in *C. albicans*, which can attach to host organs or medical devices and thus play an important role in adhesion and penetration into host tissues (4, 5). To investigate whether small molecules B and C inhibit biofilm formation, biofilm was formed by densely attaching *C. albicans* to 96-well flat-bottomed microtiter plates for 24 h at 37°C. After the biofilm was densely formed, small molecules B and C were treated to a final concentration ranging from 0.125–64 μg/mL. Surprisingly, the results showed that two molecules with MICs of 8 μg/mL and 16 μg/mL, respectively, inhibited biofilm formation at a lower concentration of 1 μg/mL (Fig. 2C). Small molecules B and C inhibit hyphal morphogenesis, which is known to be closely related to the virulence of *C. albicans* and effectively inhibit biofilm formation, the final form of this morphogenesis. This suggests that these small molecules will greatly inhibit the pathogenicity of *C. albicans.*

**Fig. 2.**
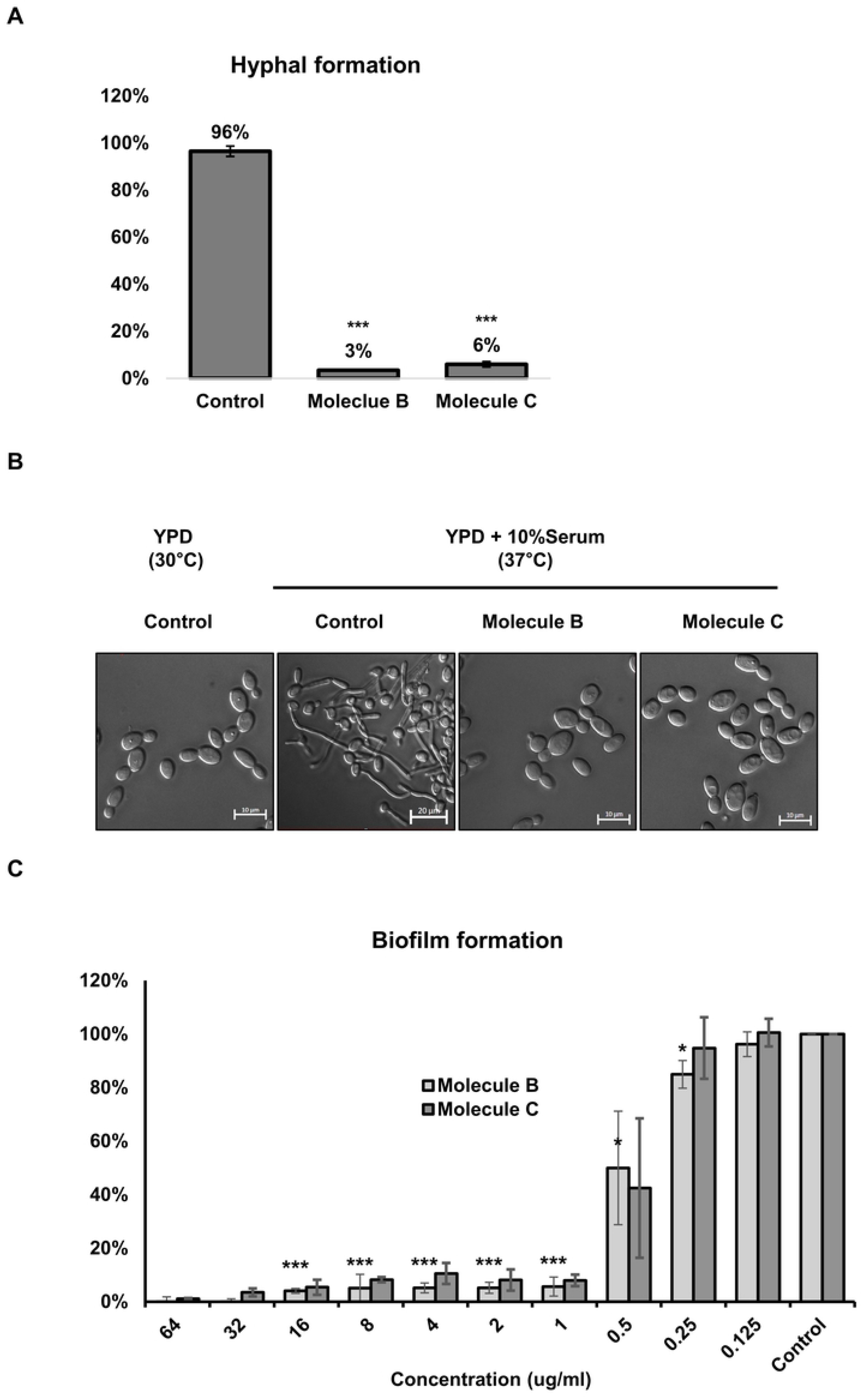
The inhibitory effect of the small molecules on morphogenesis and biofilm formation in *C. albicans.* (A) The inhibitory effects of the small molecules on *C. albicans* hyphal formation. For conditions for hyphae induction, YPD with 10% FBS was used and incubated for 2 h at 37 °C. A minimum of 300 cells were counted for each treatment to evaluate the morphogenesis inhibitory effect as a percentage. (B) To observe the morphology change using a microscope, small molecules B and C were treated at each MICs. Scale bars represent 10μm(yeast form) and 20μm(hyphae). (C) The inhibitory effect of the small molecules on *C. albicans* biofilm formation. Biofilms were formed by inoculation 2 × 10^4^ cells/mL in 200 μL cell suspension into 96-well flat-bottomed microtiter plates for 24 h at 37°C. The small molecules were added and the plates were incubated for another 16 h at 37°C. A metabolic assay based on the reduction of XTT was performed to determine the biofilm formation inhibitory concentrations. Colorimetric absorbance was measured at 495 nm in a microtiter plate reader. Each experiment was conducted in triplicate. The data represent the mean and standard deviation of three independent experiments. *; P<0.05, **; P<0.01, ***; P<0.001 (*t*-test).

### Virulence diminishes in the candidiasis murine model treated with the small molecules via oral intake and vein injection

*In vivo* experiments were conducted based on the results that the small molecules potently inhibited hyphal morphogenesis of *C. albicans.* To conduct the *in vivo* experiment, a candidiasis mouse model was established, and the survival rate and degree of Candida infection in the kidney, an organ known as a major target of *C. albicans* (19), were investigated. The survival rate of the groups receiving the small-molecule treatment through tail vein injection was 100%, but the control group died within 9 days (Fig. 3A). The groups receiving the small molecule treatment had a very low level of infection compared to the control group (Fig. 3B). In addition, we investigated whether the degree of candidiasis infection was reduced by ingestion of the small molecules dissolved in drinking water. The groups of the candidiasis mice ingesting the small molecules by autonomous intake using the small molecules dissolved in water showed very high survival rates compared to the control group (Fig. 3C). In addition, the results of the investigation on the level of Candida kidney infection showed very low infection rates compared to the control group (Fig. 3D). These results imply that the small molecule treatment potently inhibits hyphal formation in *C. albicans*, the morphological transition, and reduces pathogenicity *in vivo*.

**Fig. 3.**
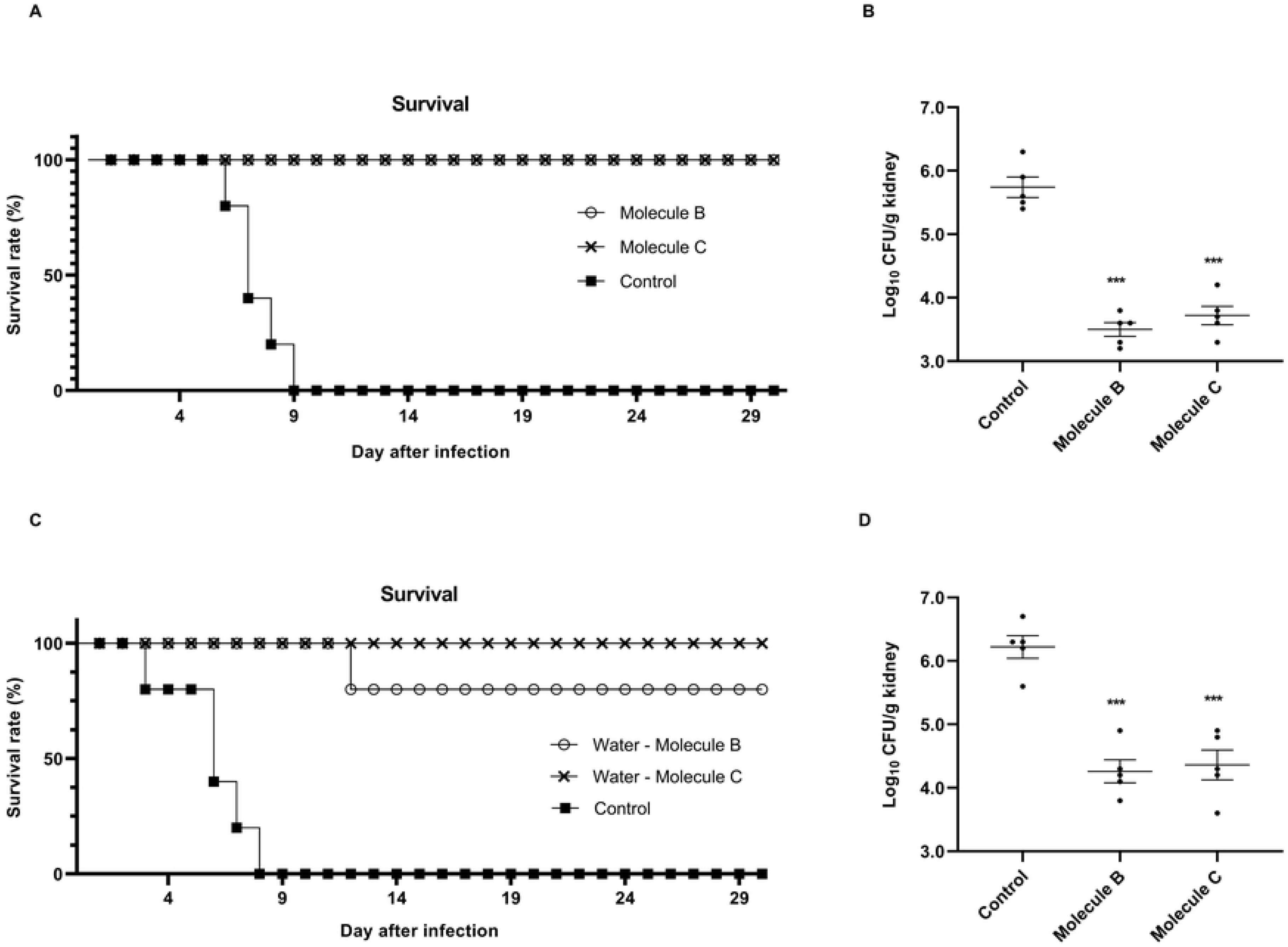
Abolishment of *C. albicans* pathogenicity by the small molecule treatment in the candidiasis murine model. The candidiasis mice model was established by inoculation of *C. albicans* 5 × 10^6^ cells through tail vein injection. (A) From the day after *C. albicans* infection to the 30th day, small molecules B and C were injected at 8 mg/kg and 16 mg/kg, respectively, compared with the weight of the mice via tail vein injection, and the survival rate was measured every day. (B) The dead mice were dissected, the kidneys were harvested, weighed, washed with PBS, chopped, spread on YPD plates, incubated at 30°C for 2 days, and then the CFUs were counted to analyze the burden of fungal infection. (C) For 1 month from the day after *C. albicans* infection, small molecules B and C were diluted to 8 mg/L and 16 mg/L, respectively, in the drinking water of mice, and the survival rate was measured every day after allowing the mice to ingest autonomously. (D) The dead mice were dissected, the kidneys were harvested, weighed, washed with PBS, chopped, spread on YPD plates, incubated at 30°C for 2 days, and then the CFUs were counted to analyze the fungal infection burden. *; P<0.05, **; P<0.01, ***; P<0.001 (*t*-test).

### Small molecules inhibit the morphogenesis of *C. albicans* by regulating the protein levels of Nrg1 and Ume6

Since HSGs are closely related to the pathogenicity of *Candida* through filamentous growth and invasive growth, we investigated whether the effect of small molecules suppressed the pathogenicity of *Candida* by lowering the level of HSGs (20, 21). As expected, expressions of the hyphal-specific genes *HWP1*(22, 23)*, ALS3*(20, 24)*, HGC1*(25), and *ECE1*(26) were suppressed under small-molecule treatment conditions (Fig. 4A). CaUme6, which upregulates HSGs, is a transcription factor that significantly affects hyphal formation (6). In addition, Nrg1 and Ume6 act as negative feedback regulators at the transcription level under filament growth conditions (21, 27). As a result of observing the change in the levels of these two proteins according to the hyphae morphology induction time (0–2 h), the level of Nrg1 protein treated with the small molecules confirmed at all times was similar or slightly higher than that at 0 h (Fig. 4B). Significantly increased levels of mRNA expression of *NRG1* were observed in the small-molecule treated cells (Fig. 4C). Ume6 protein was not detected at all times in the cells treated with the small molecules (Fig. 4D). A markedly reduced level of mRNA expression of *UME6* was observed (Fig. 4E). To investigate whether morphogenesis was inhibited under hyphal-inducing conditions by treatment with the small molecules, *tetO-UME6*, in which one allele of *UME6* is driven by a tetracycline-regulating promoter, was established and used. In the presence of the tetracycline derivative doxycycline (DOX), *UME6* is constitutively expressed and *UME6* expression is blocked in the absence of DOX (28). Contrary to our expectation that even under the constitutive expression of *UME6*, the small molecule treatment might prevent hyphal formation through direct Ume6 regulation, inhibition of hyphal formation by treatment with small molecules did not occur (Fig. S2). These results indicate that small-molecule treatment inhibits the pathogenicity of *C. albicans* through regulation of HSG by indirectly controlling the level of Ume6, which play an important role in morphogenesis.

**Fig. 4.**
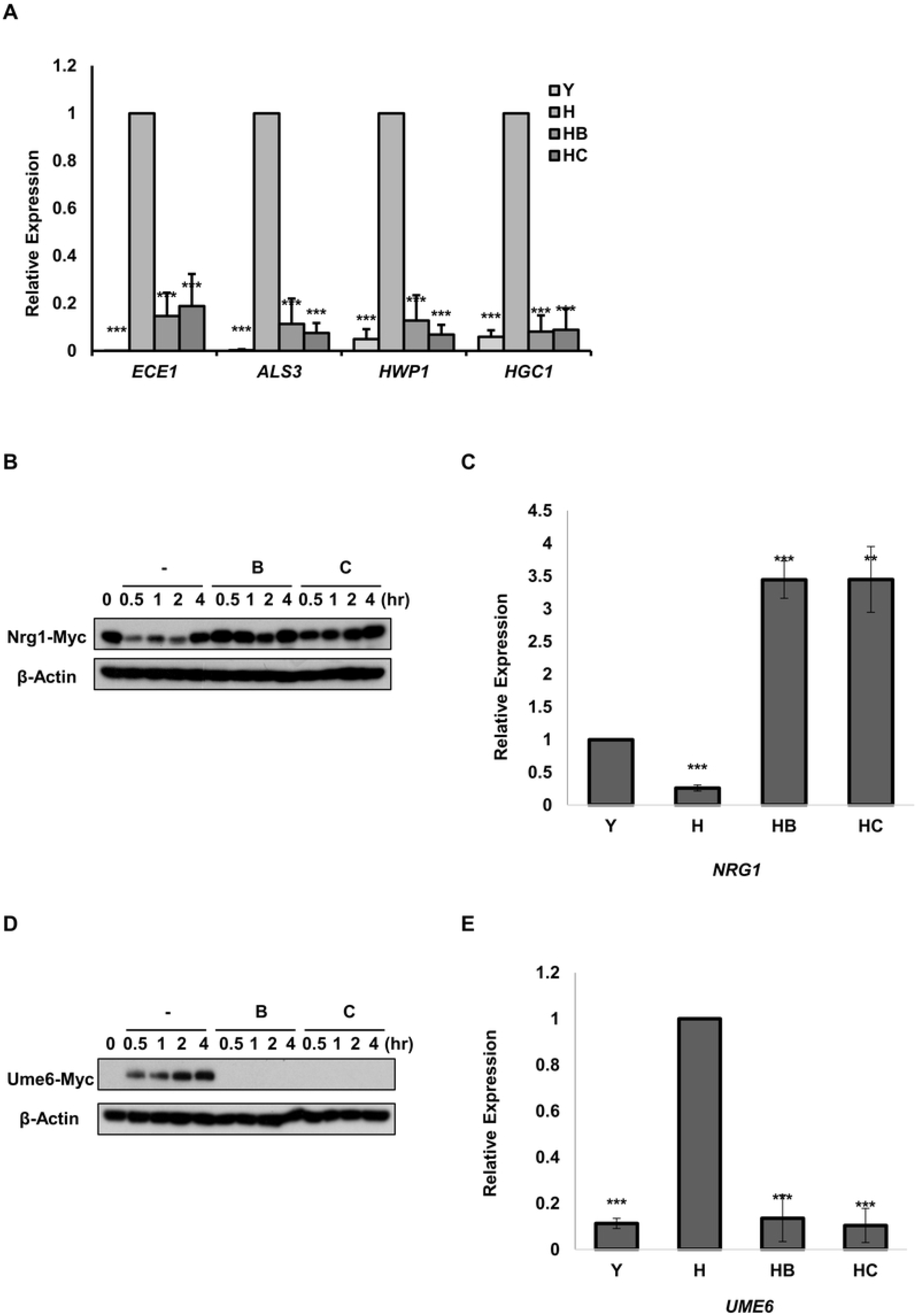
The effects of small molecules B and C on the expression of HSG, Ume6, and Nrg1 genes. (A) Total RNA of yeast cells (Y), the small molecules treated hyphae-induced cells (HB, HC), or hyphae-induced cells (H) were isolated using the TRIzol reagent method. Total RNA was measured using Nanodrop Spectrophotometer and was reverse transcribed into cDNAs using oligo-d(T) and M-MLV reverse transcriptase. Real-time PCR was performed with the specific primers listed in Table 2. (B) Immunoblotting was performed by preparing protein samples for 0–4 h, and Real-time PCR (C) was performed as previously described using the Nrg1-myc strain. (D) Immunoblotting was performed by preparing protein samples for 0– 4 h, and Real-time PCR (E) was performed as described above using the Ume6-myc strain. Western blotting was performed with an anti-myc antibody to detect Nrg1-myc and Ume6-myc. In the western blotting, β-actin was used as a loading control by detecting with the anti-β-actin antibody. Each experiment was conducted in triplicate. The data represent the mean and standard deviation of three independent experiments. *; P<0.05, **; P<0.01, ***; P<0.001 (*t*-test).

### The small molecules regulate the Ras1 and MAPK pathways

Ras signaling is known to mediate hyphal growth induction in response to a variety of environmental signals (3). Ras signaling is transmitted through the cyclic AMP (cAMP)-protein kinase A (PKA) pathway and the mitogen-activated protein (MAP) kinase pathway (29), these two established cAMP-PKA/MAPK signaling pathways are involved in the hyphal formation of *C. albicans*. The expression of known genes in these kinase cascade pathways, such as *RAS1, CYR1, TPK1, EFG1, CST20, HST7, MKC1, CPH1*, and *CEK1* was analyzed using quantitative real-time PCR. Notably, the result confirmed that the expression of *CSt20, HST7*, and *CPH1* corresponded to the kinase cascade genes related to *CEK1* (Fig. 5A), one of the MAPKs in *C. albicans*, and the expression of *RAS1* decreased in the cells treated with small molecules (Fig. 5B); another MAPK, *MKC1*, also decreased (Fig. 5C). As expected, the level of Ras1 protein in the cells treated with small molecules was very low compared to that of the untreated hyphae-formed cells (Fig. 5D). The decreased levels of Cek1 and Mkc1 were confirmed, and the levels of phosphorylated Cek1 and Mkc1, the activated forms, were also confirmed (Fig. 5E). As with the Mkc1 level, there was no change in the level of the active form, phosphorylated Mkc1, but interestingly, the levels of phosphorylated Cek1 reduced significantly in the cells treated with the small molecules (Fig. 5E). These results suggest that the treatment effect of the small molecules functions through a key switch called Ras1, and is achieved by inhibiting mainly Cek1 cascade activity in the MAPK pathway, which is well-known to be associated with the morphological transition in *C. albicans (30, 31)*.

**Fig. 5.**
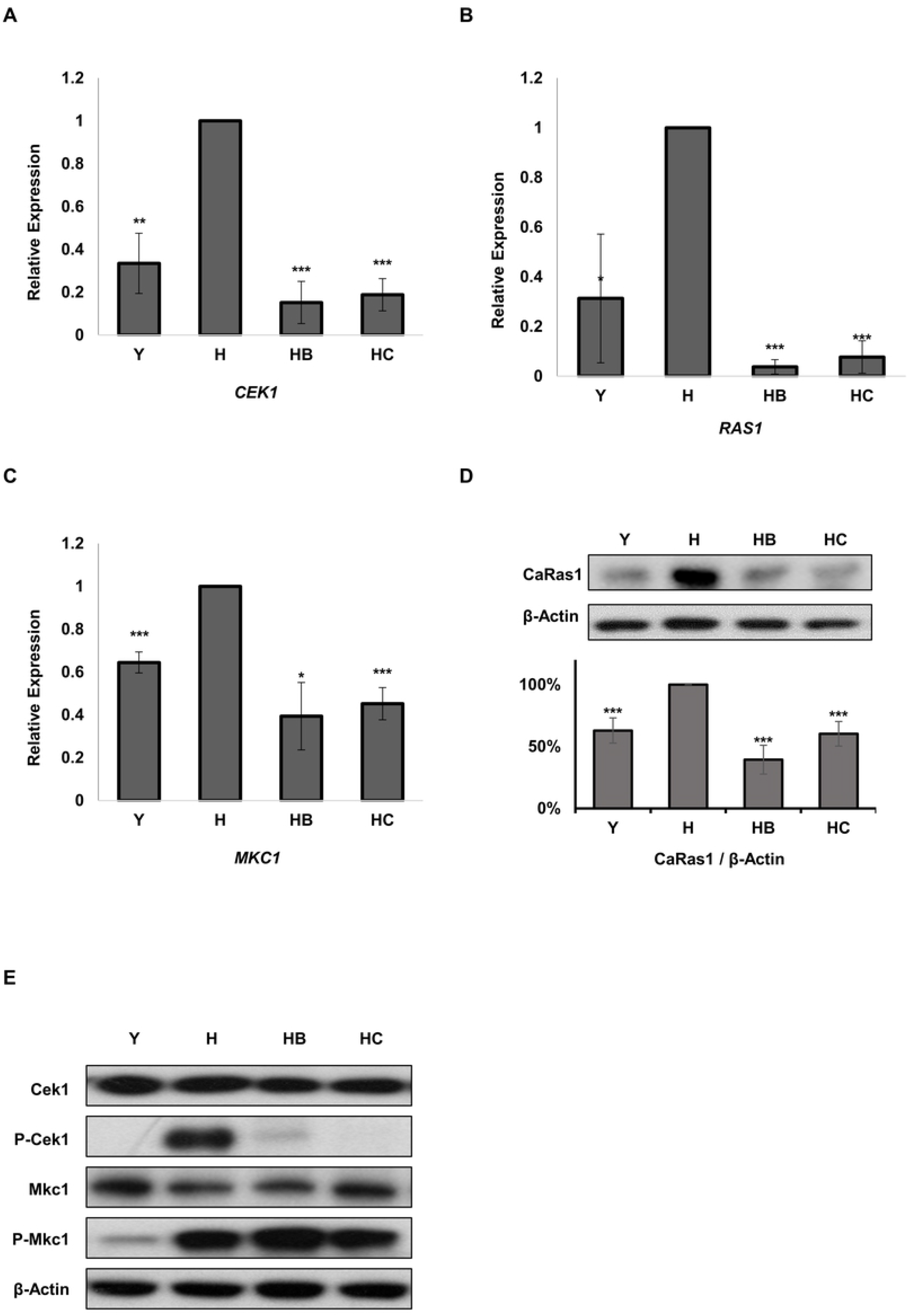
The mRNA expression and protein levels of *RAS1* and MAPK were reduced by small molecule treatment, but the activated form of P-Mkc1 did not decrease, only P-Cek1 did. Real-time PCR was performed to investigate the relative mRNA expression levels of yeast cells and small molecules treated hyphae-induced cells against hyphae-induced cells. The relative mRNA expression levels of CEK1(A), RAS1(B), and MKC1(C) were investigated with real-time PCR. (D) In the western blotting, CaRas1 was detected by anti-Ras10 antibody and β-actin was used as a loading control. The quantification graph by image J is expressed by normalizing the ratio of each CaRas1 blot to each β-actin loading control. (E) Protein samples were prepared from yeast cells, hyphae cells, and small molecules treated hyphae cells. Western blotting was performed by detecting Phospho-Mkc1 and Phospho-Cek1 with an anti-phospho-p44/42 antibody. Mkc1 and Cek1 were detected by anti-p44/42 antibody and β-actin was used as a loading control. Each experiment was conducted in triplicate. The data represent the mean and standard deviation of three independent experiments. *; P<0.05, **; P<0.01, ***; P<0.001 (*t*-test).

## Discussion

In this study, we demonstrated that two small molecules inhibit virulence in *C. albicans* via regulation of the Ras1/MAPK pathway. These two small molecules that inhibit *C. albicans* growth and are not cytotoxic in mammalian cells were selected from 32 small molecules containing the piperazinyl carbazole structure through screening. In spite of some studies have already been reported in which several small molecules containing a carbazole structure exhibit growth inhibitory activity against bacteria and *Candida* species (15–17), this study first showed the mechanism by which these two small molecules inhibit the pathogenicity in *C. albicans* through inhibition of the Ras1/MAPK pathway.

### The small molecules reduced the pathogenicity of *C. albicans* by inhibiting the morphological transition

Numerous studies have suggested that morphogenesis in *C. albicans* is closely related to pathogenicity (1) and the development of these hyphae leads to attachment to host tissues or medical devices and forms biofilms (4). In *in vitro* experiments, the small molecules inhibited hyphal formation. In addition, the biofilm formation was also suppressed (Fig. 2C). The proteins Als2, Als3, and Als4 were investigated to promote *C. albicans* adhesion to host cells, and E-cadherin in epithelial cells or N-cadherin in endothelial cells were studied as binding targets for Als3 (32). Thus, it is considered that the effect of inhibiting biofilm formation even at a concentration lower than that of the MICs of the small molecules is a result of the suppression of the expression level of *ALS3* (Fig. 4A). The survival rates of mice were very high when the small molecules were administered, and the degree of kidney infection was also very low (Fig. 3). These results confirmed that the two small molecules reduced the pathogenicity of *C. albicans* by inhibiting morphological transition.

### The inhibition of morphological transition by treatment with the small molecule is associated with inhibition of the Ras1/MAPK pathway

In this study, we found that *UME6* and Ume6 expression levels were inhibited by treatment with the small molecules. Ume6 in *C. albicans* is an important transcriptional factor for HSGs (6) and the small molecules repressed HSGs such as *HWP1, ALS3, HGC1*, and *ECE1* (Fig. 4A). In addition, both Nrg1 and Ume6 act as negative feedback at the transcription level under filamentous growth conditions (21, 27), and our study also confirmed that the expression levels of NRG1 and Nrg1 increased under small-molecule treatment conditions. From these results, treatment with the small molecules under filamentous growth conditions reduced morphogenesis, a pathogenic factor, through the inhibition of the expression level of HSGs via the regulation of the key transcriptional factors Nrg1 and Ume6 (Fig. 4B-4E). Ras1, a key element of the signaling pathway upstream of hyphal formation in *C. albicans*, controls the induction of morphogenesis in response to multiple stimuli, including serum and elevated temperature, and stimulates both the PKA and MAPK pathways (2, 3). According to the qRT-PCR results of this study, the expression levels of genes encoding protein factors related to MAPK signaling and *RAS1* were significantly reduced when treated with small molecules (Fig. 5B and 5D). In *C. albicans*, four MAPKs mediate signaling: Cek1, Cek2, Hog1, and Mkc1. Hog1 is involved in adaptation to osmotic and oxidative stress (33). Mkc1 mediates signals in cell wall construction (34) and invasive growth (35). Similar to Mkc1, Cek1 regulates invasive growth (30) and cell wall construction (31), and Cek2 is also involved in mating (36). As described in the preceding paragraphs, Cek1 and Mkc1 were found to be closely related to pathogenicity in *C. albicans* because they mediate signaling for cell wall reconstruction and invasion into host cells, which are deeply related to hyphal growth (3). In this study, it was confirmed that the phosphorylation of Cek1 and Mkc1 increased under conditions that induce hyphal formation in the serum. Although the transcriptional levels of *MKC1* and *CEK1* decreased upon treatment with small molecules, the decrease in phosphorylation was confirmed only in Cek1 (Fig. 5A, 5C, and 5E). This supports our hypothesis that among several MAPK pathways, only the Cek1 signaling pathway was inhibited by treatment with small molecules. A study on hyphal developmental signaling in *C. albicans* has shown that Cph1, a downstream transcriptional factor of Cek1 in the MAPK pathway and Ras1, an upstream signal transducer, directly regulates the transcriptional level of *UME6*, a major target of many transcriptional regulators. Consequently, Ume6 regulates the expression of HSGs (37). Consistently, our results confirmed that the phosphorylation of Cek1 was reduced by treatment with the small molecules under hyphal formation-inducing conditions. The decreased transcriptional level of *CPH1*, which is downstream of Cek1, and thus the decreased transcriptional level of *UME6* and the expression level of Ume6 were confirmed (Fig. 4 and S3).

### A model of how the small molecules inhibit virulence in *C. albicans*

Our results from the model in Figure 6 shows how the small molecules regulate filamentous growth in *C. albicans*. The small molecules regulate ‘*RAS1* and *CEK1’* mediated MAPK pathway (*STE11-HST7-CEK1*) which is associated with hyphal cell wall construction and invasive growth. We looked for other targets that would be affected by Ras1 activation and consequently confirmed that the expression levels of genes involved in the MAPK pathway were effectively reduced (Fig. S3). In the CEK1-mediated pathway, MAPK kinase kinase (MAPKKK) *STE11* is reduced following the downregulation of Ras1 (30). In addition, MAPK kinase (MAPKK) *HST7* is reduced (30), and finally, the transcriptional factor *CPH1*, downstream of the *CEK1* pathway (38), is reduced. *CPH1* is involved in the regulation of HSG expression either directly or through *UME6*, a major target of many transcriptional regulators (39). In addition, a significant increase in *NRG1* expression was confirmed in the small molecule-treated environment. *NRG1* directly regulates HSG expression or indirectly through the inhibition of *UME6* expression (21, 27, 39).

**Fig. 6.**
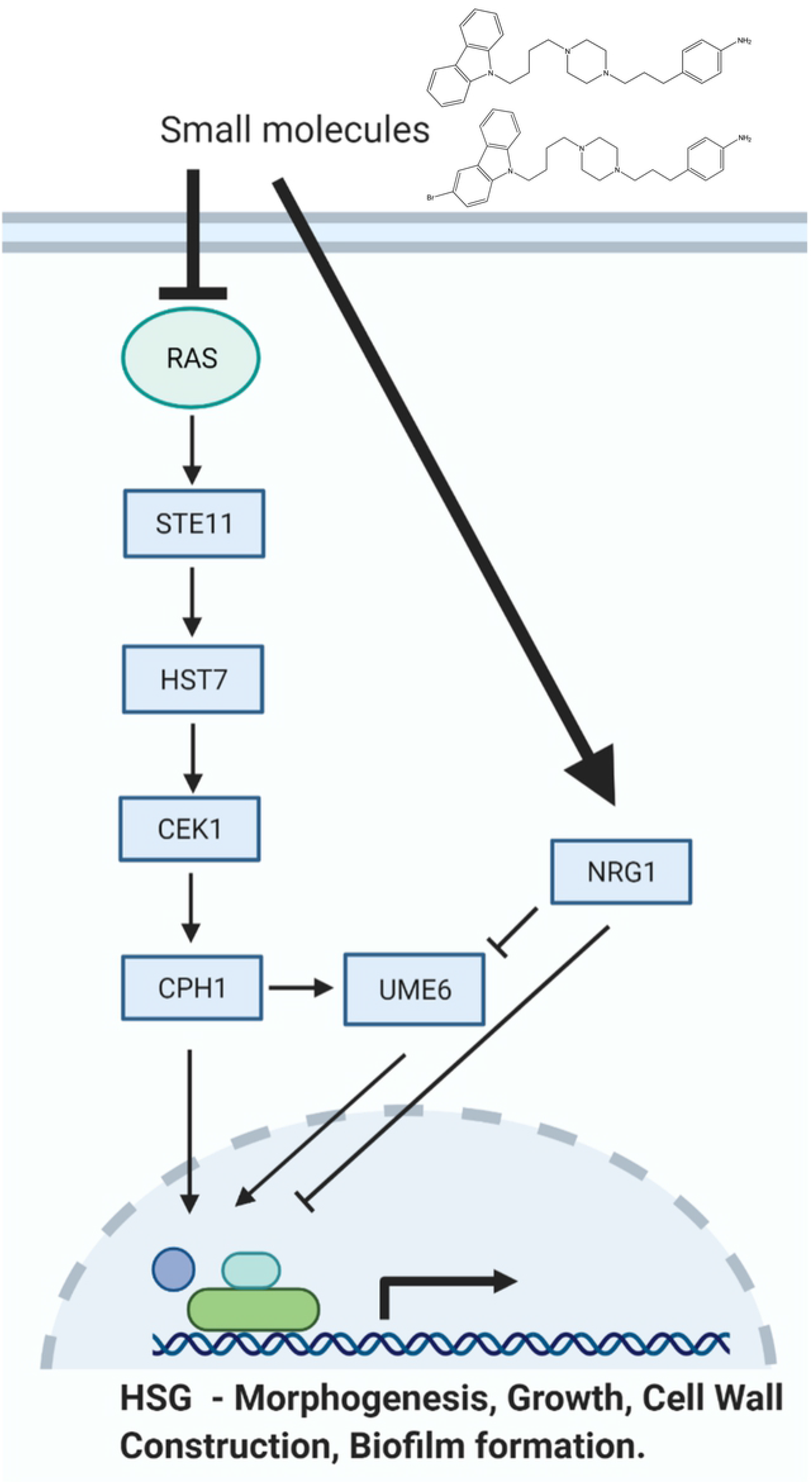
Putative model of virulence inhibition in *C. albicans* by the small molecules. The inhibitory effects of the small molecules on upstream activator signaling proteins are shown. The small molecules inhibit Ras1 and affect the PAK kinase Cst20, thus affecting the Cek1 MAP kinase cascade, resulting in a decrease in the transcription factor Cph1. Ume6, which is regulated by the signal of Cph1, is also regulated in a way that decreases as the amount of Nrg1 increases.

In this study, we introduced novel small molecules that reduce virulence by inhibiting morphogenesis, which is a pathogenic factor of *Candida* via the Ras1/MAPK pathway. In particular, it could be a promising treatment for patients suffering from *Candida* diseases, as can be seen from the results of experiments on strains that are resistant to existing therapeutic compounds. Our findings are supported by these small molecules as promising compounds for the development of new antifungal agents for the treatment of systemic candidiasis and possibly for other fungal diseases.

## Materials and Methods

### Strains, culture media, and chemicals

*C. albicans* strains, including FLC-resistant and echinocandin-resistant strains, primers, and small molecules used in this study are listed in Tables 1, 2, and 3, respectively. To generate the Ume6-myc and Nrg1-myc strains, PCR-based gene disruption was performed using HIS1 and 9myc-NAT1 cassettes. The primers for amplification of the cassettes (HIS1 and 9myc-NAT1) are listed in Table 2. To generate the UME6-9myc and NRG1-9myc strains, pFA6a-9myc-NAT1 was used for UME6-9myc-NAT1 and NRG1-9myc-NAT1 cassettes. These DNA cassettes were transformed into the *ume6::HIS1/*UME6 and *nrg1::HIS1/* NRG1 strains. To confirm strain generation, genomic DNA was isolated, and PCR and western blotting were performed with specific primers and antibodies. The cells were cultivated at 30°C in YPD medium (1% yeast extract, 2% peptone, and 2% dextrose) or for hyphal morphogenesis induction, and were incubated in YPD medium containing 10% fetal bovine serum (FBS) at 37°C with shaking. A synthetic complete medium (0.76% yeast nitrogen base without amino acids and 2% dextrose) supplemented with the appropriate auxotrophic requirements and uridine (50 μg/mL), was used to select positive transformants. Their susceptibilities were determined according to the Clinical and Laboratory Standards Institute (CLSI) document M27-A3 (18) and RPMI 1640 (pH 7.0) was used as the liquid medium for diluting the drugs and strains. *S. cerevisiae* BY4741 cells were grown at 30°C in YPD medium, as previously described (40, 41). All stock solutions of the chemicals were dissolved in sterile distilled water or methanol at a final concentration of 20 mg/mL. All stock solutions were stored at −20°C until use.

**Table 1.** *Candida albicans* strains used in this study.

| **Strain** | **Relevant genotype** | **Source** |
| --- | --- | --- |
| **SC5314** | Clinical isolate from London Mycological Reference Laboratory | (48) |
| **BWP17** | *ura3::imm434/ura3::imm434 his1::hisG/his1::hisG arg4::hisG/arg4::hisG* | (49) |
| **Ume6-myc** | As BWP17 except for *ume6::HIS1/UME6-9myc-NAT1* | In this study |
| **Nrg1-myc** | As BWP17 except for *nrg1::HIS1/NRG1-9myc-NAT1* | In this study |
| **tetO-UME6** | As BWP17 except for *ADH1/adh1::P<sub>tet</sub>-UME6* | In this study |
| **12-99** | Azole resistant strain | (50) |
| **89** | Echinocandin resistant strain | (51) |
| **177** | Echinocandin resistant strain | (51) |

**Table 2.**
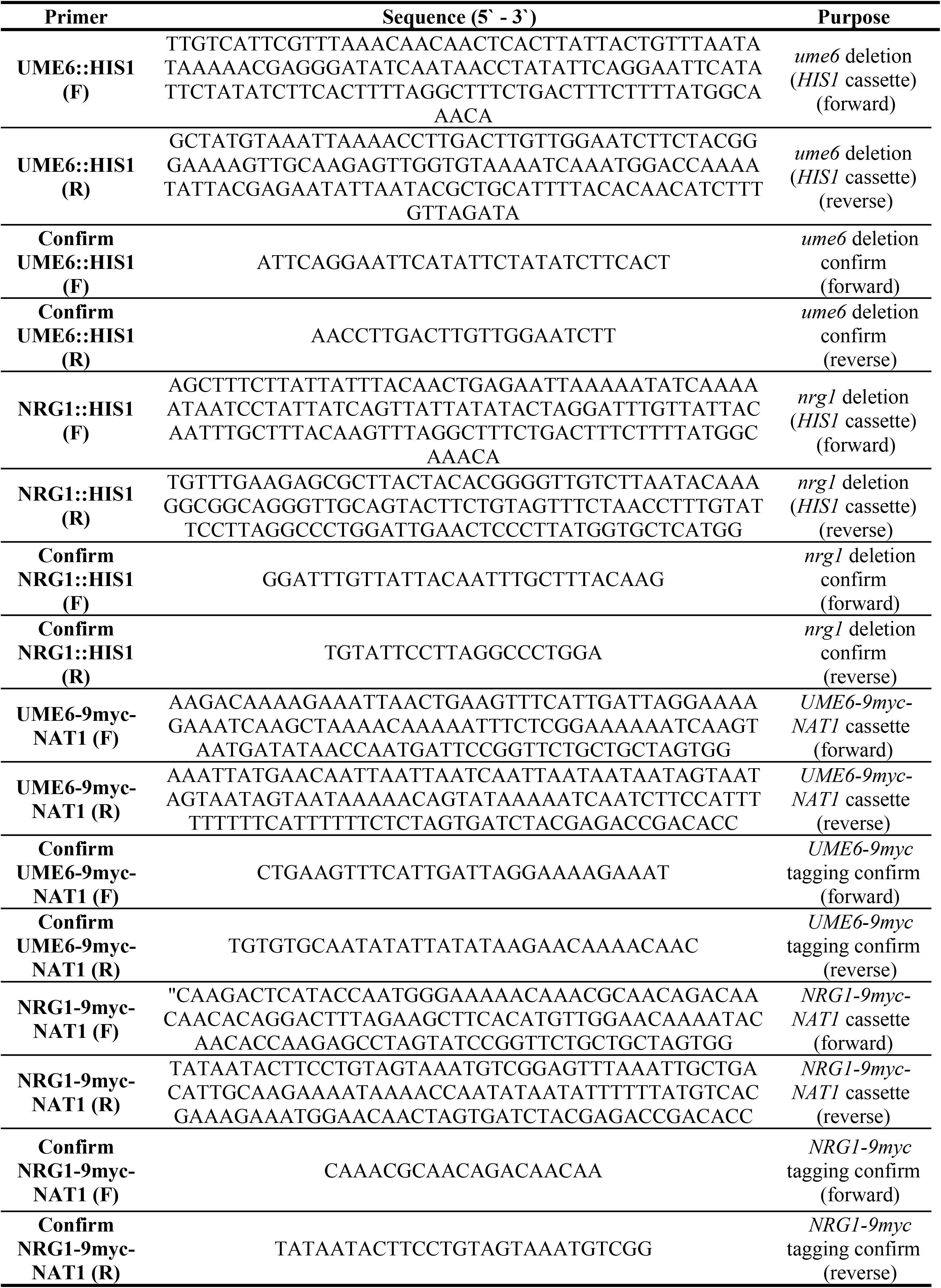
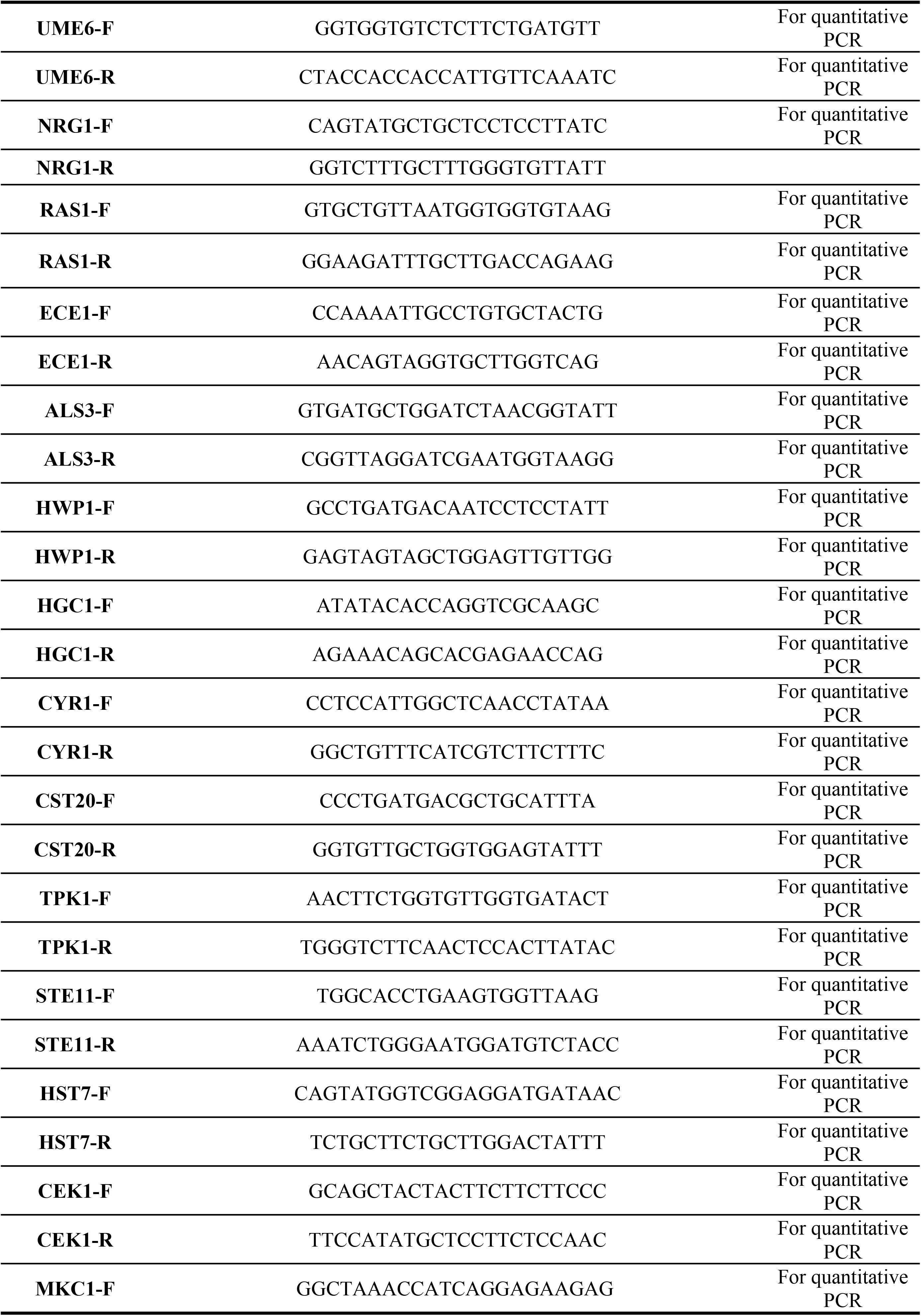
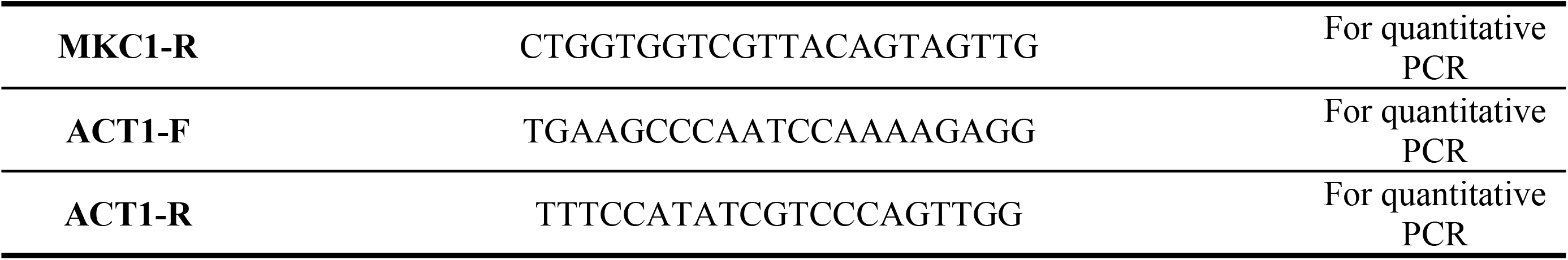
Oligonucleotides used in this study.

**Table 3.** MICs of the small molecules in *C. albicans*.

|  | Fluconazole | Caspofungin | Molecule B | Molecule C |
| --- | --- | --- | --- | --- |
| **SC5314<sup>a</sup>** | 1 | 1 | 8 | 16 |
| **12-99<sup>b</sup>** | >128 | 1 | 8 | 16 |
| **89<sup>c</sup>** | >64 | 4 | 8 | 16 |
| **177<sup>c</sup>** | 1 | 4 | 8 | 16 |

\*All units indicated are µg/mL. <sup>a</sup> Wild type *Candida albicans* strain. <sup>b</sup> Fluconazole resistant *Candida albicans* strain (50). <sup>c</sup> Echinocandin resistant *Candida albicans* strain (51).

### Hyphae formation

YPD medium containing 10% FBS at 37°C was used to test the effect of small molecules on hyphal formation. *C. albicans* (2 × 10^6^ cells/mL) was inoculated in hyphal formation-inducing media containing MICs of small molecules and incubated at 37°C with shaking for 2 h. The morphology of the cells was photographed using a Zeiss microscope (Carl Zeiss).

### Antifungal susceptibility test

The minimum inhibitory concentrations (MICs) of the small molecules against *C. albicans* were determined using the broth microdilution method as described by the CLSI guidelines (18). The tests were performed in 96-well flat-bottomed microtiter plates. The final concentration of the cell suspension in RPMI 1640 medium was 2 × 10^4^ cells/mL, and the final concentration of the small molecules ranged from 0.125–64 μg/mL. All the wells were filled with RPMI 1640 to a final volume of 200 μL. The plates were incubated at 37°C for 24 h. Growth inhibition was determined by measuring the optical density at 595 nm using a microplate reader. MIC was defined as the lowest concentration of small molecules that inhibited the growth of cells by 95% compared to the control.

### Determination of minimum inhibitory concentrations of C. albicans biofilm formation

Biofilms were formed by inoculation of 2 × 10^4^ cells/mL in 200 μL cell suspension into 96-well flat-bottomed microtiter plates for 24 h at 37°C. Each well was washed with 200 μL PBS three times to remove planktonic cells. The small molecules were added at a final concentration ranging from 0.125–64 μg/mL and the plates were incubated for another 16 h at 37°C. A metabolic assay based on the reduction of XTT (sodium 3’-[1-(phenylaminocarbonyl)-3,4-tetrazolium]-bis (4-methoxy6-nitro) benzene sulfonic acid hydrate) was performed to determine the biofilm formation inhibitory concentrations. It referred to the lowest concentrations, where there was a 95% reduction in the XTT-colorimetric readings compared with the control. Colorimetric absorbance was measured at 495 nm using a microtiter plate reader.

### Cell cytotoxicity test

The cytotoxicity of the small molecules was determined in HeLa cells. HeLa cells were maintained as previously described (42, 43) in Dulbecco’s modified Eagle’s medium (DMEM) supplemented with 10% FBS (Invitrogen) and grown at 37°C in a humidified atmosphere of 5% CO_2_. Viability was measured using the MTS assay. Each well was inoculated with 10^4^ HeLa cells and incubated at 37°C for 24 h in 96-well flat-bottom tissue culture plates using DMEM containing 10% FBS. The small molecules were added at final concentrations ranging from 0.125–64 μg/mL and incubated at 37°C for 16 h. The MTS solution with PMS was added, and the cells were incubated for 10 min. The absorbance at 490 nm was measured using a microplate reader.

### Real-time PCR

Total RNA samples of yeast or hyphal-formed cells were isolated using the TRIzol reagent method. Real-time PCR was conducted as described previously (42, 44–46). In brief, total RNA was measured using a Nanodrop 1000 Spectrophotometer (Thermo Scientific, Rockford, IL, USA) and was reverse transcribed into cDNAs using oligo-d(T) and M-MLV reverse transcriptase (Promega). Real-time PCR was performed with the specific primers listed in Table 2 using the LightCycler 480 (Roche, Manheim, Germany).

### Immunoblotting

Western blotting was performed as described previously (40, 46, 47), with slight modification. Briefly, yeast or hyphal formed cells were harvested and washed with ice-cold PBS. To prepare whole cell lysates, the cells were lysed with ice-cold buffer comprised of 50 mM of Tris-HCl (pH 7.5), 150 mM of NaCl, 5 mM of EDTA, 10% of glycerol, 0.2% of Nonidet P-40, 1 mM of dithiothreitol, protease inhibitors (1 mM of phenylmethylsulfonyl fluoride, 1 μg ml–1 of aprotinin, pepstatin A, and leupeptin), 0.1 mM of sodium orthovanadate, 20 μM of sodium glycerophosphate, 20 μM of para-nitrophenyl phosphate, and 20 μM of sodium fluoride. The cells were lysed using glass beads (Sigma-Aldrich) in a Fast prep-24 device (MP Biomedical). The proteins were resolved on 10–12% SDS-PAGE and analyzed using immunoblotting on PVDF membranes according to standard procedures. For the detection of Ras1, Ume6-myc, Nrg1-myc, Cek1, Mkc1, phosphorylated Cek1, and phosphorylated Mkc1, the primary antibody (1:3000 dilution for Ras1(05-516 Merck) and 1:1000 dilution for c-Myc(c-Myc(9E10) Santacruz) and 1:1000 dilution for both Cek1/Mkc1(9102S Cell Signaling) and 1:1500 dilution for both phospho-Cek1/phosphor-Mkc1(4307T Cell Signaling)) were used. For the loading control, β-actin antibody (ab8224 Abcam) was used at 1:1000 dilution. The membranes were blocked using Tris-buffered saline containing 0.1% Tween (TBST) containing 5% skimmed milk. BM Chemiluminescence western blotting Substrate kit (POD) was used to develop the blot.

### Antifungal activity test in the murine candidiasis model

Overnight-cultured *C. albicans* strains were pelleted and washed three times with 1 mL of sterile PBS. Next, 1.0 × 10^6^ cells were resuspended in 200 μL PBS and injected into 5-week-old BALB/c (female) mice via lateral tail veins. After the establishment of the murine model, mice (n= 5/group) were treated by tail vein injection with small molecules (8 mg/kg or 16 mg/kg) or PBS every day from the second day. To investigate the voluntary intake effect, the mice were treated with small molecules (8 μg/mL or 16 μg/mL) or normal water every day. The dead mice were dissected, the kidneys were harvested, weighed, washed with PBS, chopped, spread on YPD plates, incubated at 30°C for 2 days, and then the CFUs were counted to analyze the fungal infection burden. The mice were monitored for 1 month, and Kaplan– Meier survival curves were plotted.

### Ethics Statement

This study was performed in accordance with the guidelines of the Institutional Animal Care and Use Committee of Korea University. The protocol and experiments were approved by the Institutional Animal Care and Use Committee of Korea University. The permit number is KUIACUC-2019-0061. All the mice used in the experiments were sacrificed with minimal suffering.

## Acknowledgments

This work was supported in part by NRF-2020R1A2C2100803

Y.P. designed the study, performed all experiments, analyzed the data and wrote and edited the manuscript and figures. Y.P. carried out all the experiments including the animal experiments. J.K., J.S. and H.L. designed and supervised the fabrication of small molecules. Y.P., H.K and J.K. contributed to the interpretation of the results. J.K. designed and supervised the whole work and wrote the manuscript. All authors discussed the results and contributed to the final manuscript.

---

**Fig S1. The cytotoxicity of small molecules B and C in *C. albicans*, *S. cerevisiae*, and mammalian cell.** (A) The cytotoxicity of small molecules B and C on *C. albicans* SC5314 strain cells (2 × 10^4^ cells/mL) were grown with small molecules in RPMI1640 medium with 0.165M MOPS for 24 h at 37°C according to the CLSI guidelines (18). The growth inhibition was determined by measuring the optical density at 595 nm using a microplate reader. (B) The cytotoxicity of small molecules B and C on *S. cerevisiae*. BY4741 strain cells (2 × 10^4^ cells/mL) were grown with small molecules in YPD medium 24 h at 30°C. The growth inhibition was determined by measuring the optical density at 595 nm using a microplate reader. (C) The cytotoxicity of small molecules B and C on mammalian cells. Viability was measured based on the MTS assay. Each well was inoculated with HeLa cells (10^5^ cells/mL). After incubation at 37°C for 24 h in DMEM medium containing 10% fetal bovine serum. The final concentrations of the small molecules (ranging from 0.125–64 μg/mL) were added and incubated at 37°C for 16 h. The absorbance at 490 nm was measured with a microtiter plate reader. Each experiment was conducted in triplicate. The data represent the mean and standard deviation of three independent experiments. *; P<0.05, **; P<0.01, ***; P<0.001 (*t*-test).

**Fig S2. Morphology of tetO-*UME6* strain treated with small molecules B and C with and without doxycycline.** For overnight cultivation at 30°C in YPD, the cells were inoculated with or without dox in a fresh YPD medium containing or without the small molecules and incubated with shaking at 30°C for 2 h. The morphology of the cells was photographed with a microscope. Scale bars represent 10μm(yeast form) and 20μm(hyphae).

**Fig S3. Quantitative RT-PCR analysis of MAPK cascade-related genes in *C. albicans*.** Yeast cells were cultivated at 30°C in a YPD medium and morphogenesis-induced cells were incubated with or without treatment with small molecules at 37°C in a YPD medium containing 10% FBS. The expression of RAS1 and MAPK cascade-related genes was investigated using qRT-PCR and the primers listed in Table 2. Data information: Each experiment was conducted in triplicate. The data represent the mean and standard deviation of three independent experiments. *; P<0.05, **; P<0.01, ***; P<0.001 (*t*-test).

